# SARS-CoV-2 genome evolution exposes early human adaptations

**DOI:** 10.1101/2020.05.26.117069

**Authors:** Erik S. Wright, Seema S. Lakdawala, Vaughn S. Cooper

## Abstract

The set of mutations observed at the outset of the SARS-CoV-2 pandemic may illuminate how the virus will adapt to humans as it continues to spread. Viruses are expected to quickly acquire beneficial mutations upon jumping to a new host species. Advantageous nucleotide substitutions can be identified by their parallel occurrence in multiple independent lineages and are likely to result in changes to protein sequences. Here we show that SARS-CoV-2 is acquiring mutations more slowly than expected for neutral evolution, suggesting purifying selection is the dominant mode of evolution during the initial phase of the pandemic. However, several parallel mutations arose in multiple independent lineages and may provide a fitness advantage over the ancestral genome. We propose plausible reasons for several of the most frequent mutations. The absence of mutations in other genome regions suggests essential components of SARS-CoV-2 that could be the target of drug development. Overall this study provides genomic insights into how SARS-CoV-2 has adapted and will continue to adapt to humans.

**SUMMARY:** In this study we sought signals of evolution to identify how the SARS-CoV-2 genome has adapted at the outset of the COVID-19 pandemic. We find that the genome is largely undergoing purifying selection that maintains its ancestral sequence. However, we identified multiple positions on the genome that appear to confer an adaptive advantage based on their repeated evolution in independent lineages. This information indicates how SARS-CoV-2 will evolve as it diversifies in an increasing number of hosts.

## INTRODUCTION

A better understanding of the origin and evolution of the SARS-CoV-2 pandemic may help to mitigate disease outbreaks. Initial genome comparisons point toward a proximal origin in horseshoe bats (1), with possible intermediate hosts including pangolins, cats, and ferrets (2–4). Differences between host species are believed to present many barriers to host switching, likely resulting in a virus that is initially maladapted to a new host (5, 6). Viruses are expected to quickly adapt to a new species via mutations that increase transmissibility and decrease the serial interval (7). Yet, relatively little is known about the mode and tempo of evolution at the start of many epidemics, including the SARS-CoV-2 pandemic. Owing to the relatively high mutation rates of RNA viruses, comparison of genome sequences may reveal a wealth of information even at early stages of an outbreak (8).

Previous studies suggest that new environments provide the opportunity to develop a large number of beneficial mutations, which may accrue quickly due to natural selection (9). High rates of adaptation have been observed for viruses propagated in cell lines belonging to new host species (10). The possibility of an adaptive advantage over the ancestor gives rise to three outcomes: (i) increasing frequencies of beneficial mutations, (ii) parallel evolution, where the same mutation rises to detectable frequencies in different lineages, and (iii) positive selection, where the number of non-synonymous changes exceeds the number of synonymous changes in protein coding regions of the genome. Mutations can rise in frequency for reasons other than positive selection, such as arising by chance near the outset of a pandemic or after undergoing a bottleneck when invading a new population of susceptible hosts. It is also challenging to accurately determine mutant frequencies due to an uneven sampling of genomes. Therefore, repeated parallel evolution is a clearer signal of adaptation than changes in frequency. Evidence of parallel evolution also enables identification of beneficial mutations before they have had time to substantially rise in frequency, which is particularly useful at the beginning of a pandemic.

At the other end of the spectrum is the mode and tempo of evolution that is expected when an organism is well-poised to enter a new environment or has spent a long time adapting to a specific context. In such cases we anticipate evolution to occur slowly because further mutation would not provide an advantage over recent ancestors. This would appear in the genome as neutral or purifying selection, where the number of non-synonymous changes does not exceed the number of synonymous changes at protein coding sites. Invasion into an environment where an organism was already well-suited would cause a bottleneck event that results in a homogenous population and a low rate of divergence due to purifying, or negative selection. Therefore, the frequency and type of mutations at the outset of a pandemic can provide insight into the ways in which a pathogen is initially well-adapted or maladapted, potentially informing the development of therapies that target its weaknesses and avoid resistance evolution.

## RESULTS AND DISCUSSION

### SARS-CoV-2 is undergoing purifying selection

Based on previous studies from similar coronaviruses, SARS-CoV-2 is anticipated to have a relatively low per-base pair mutation rate for RNA viruses, resulting in approximately 1 mutation per genome replication (11–13). We estimated an average viral replication time of 6 hours based on SARS-CoV-2 titers and the known replication times of similar viruses (14).

Therefore, we would expect mutations to accumulate on the order of ~4 per day under neutral evolution and potentially faster under positive selection. In contrast, we observe the number of nucleotide substitutions increasing at a rate of approximately 0.062 per day (Fig. 1). This equates to 8 × 10^-4^ per site per year, which is similar in magnitude to other RNA viruses and indistinguishable from that of SARS-CoV (15). This result implies that purifying selection dominated during the early stages of the SARS-CoV-2 pandemic. However, it does not rule out the possibility that some sites are under positive selection even if most remain under negative selection.

**Figure 1.**
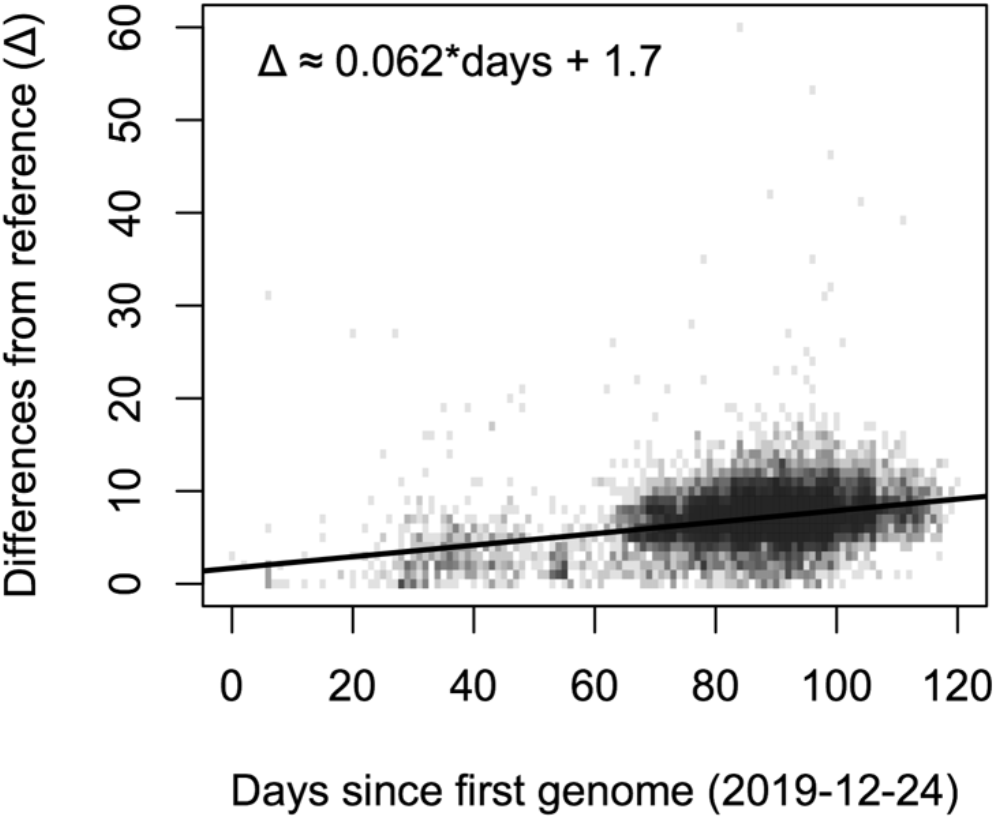
Rate of acquiring substitutions by SARS-CoV-2. The number of genomes with a given number of nucleotide substitutions relative to the reference genome (NC_045512.2) are shown since the first collected genome. The number of substitutions has increased at an average rate of approximately 0.062 substitutions per day (line of best fit), which is substantially lower than expected from a neutral model of evolution (~1 per day) and indicative of purifying selection.

### Mutational and selection biases across the SARS-CoV-2 genome

Beneficial mutations can be readily identified in laboratory evolution experiments through parallel mutations that arise in multiple independent replicates (16, 17), also known as homoplasies. We applied this intuition to search for beneficial mutations in the SARS-CoV-2 genome as it diversifies across many hosts. As shown in Figure 2, we observed 6,028 positions with substitutions out 29,903 nucleotides in the genome (Fig. 2a). Of these, 2,070 positions had more than one independent substitution, 1,858 of which were located in coding regions. Substitutions displayed a strong bias toward guanine (G) and cytosine (C) being replaced with uracil (U) in the genome (Fig. S1). U replacing C was 2.5-fold more common than the reverse, and U replacing G was 6.4-fold more common than the reverse. C to U transitions accounted for 31% of substitutions, and may result from effects of the APOBEC3G gene causing deamination of C to U (18). G to U substitutions represented 43% of transversions, although the cause of their relatively high frequency was unknown.

**Figure 2.**
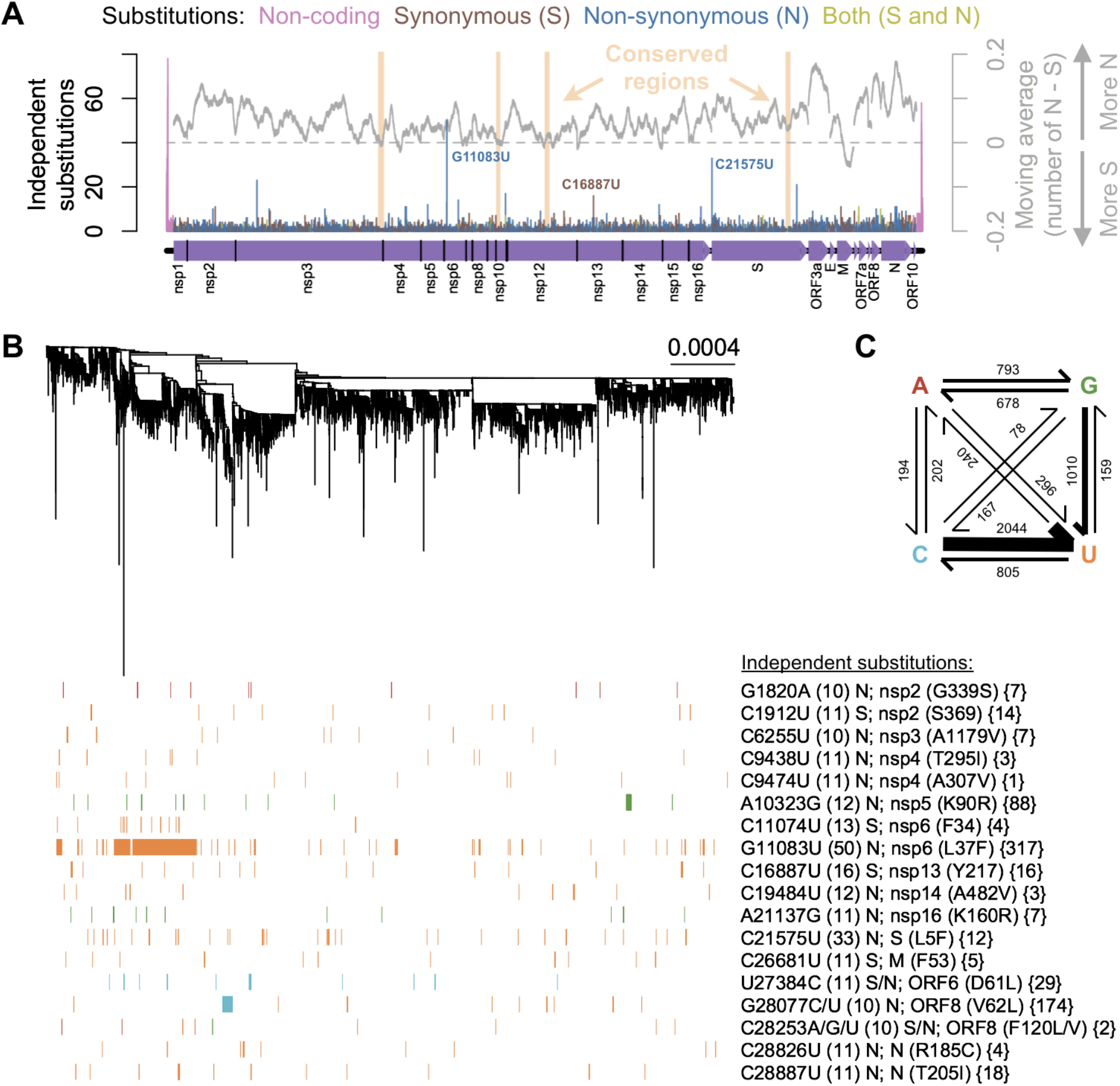
Mutational biases in the SARS-CoV-2 genome. (A) Independent substitutions were unevenly distributed along the genome, with a high concentration of parallel mutations near the genome termini and a paucity of substitutions in some coding regions. Long (≥ 100 nucleotide) conserved regions with few substitutions indicate where the genome is more constrained and could be the focus of drug targets that avoid resistance development. Some proteins (e.g., M) displayed a bias toward synonymous substitutions, suggesting the dominance of purifying selection purging changes to the protein sequence. (B) The pattern of parallel substitutions with 10 or more independent occurrences (number in parentheses) across a rooted phylogenetic tree built from SARS-CoV-2 genomes. Of these 18 substitutions, 14 change the protein sequence (N) and four are synonymous (S). The size of the largest clade associated with each mutation is shown in braces. (C) The matrix of substitutions shows a bias toward cytosine or guanine to uracil mutations.

Non-synonymous substitutions represented 66% of substitutions in coding regions. Based on the frequency of codons and observed substitution rates, we estimated that non-synonymous substitutions would represent 71% of substitutions without selection against changes to the protein sequence. Therefore, the slight bias toward non-synonymous substitutions is lower than expected and consistent with purifying selection being the dominant mode of evolution. However, the skew toward non-synonymous substitutions varied across the genome (Fig. 2a), reaching a peak within the gene coding for the nucleocapsid protein N that protects the viral RNA. In contrast, genes with a bias toward synonymous substitutions, such as the membrane protein M, indicate their protein sequence is relatively constrained.

We observed several conserved regions that displayed a relative lack of mutations (Fig. 2a), including the C-terminus of nsp3 (1,880 – 1,959), the N-terminus of nsp10 (2–59), a central region within the RNA polymerase nsp12 (504 – 570), and a region within the spike protein S (976 – 1,041). The conserved region within nsp12 overlaps with the entry tunnel for the RNA template (19) and the predicted binding sites of many antivirals (20, 21). The conserved region within S encompasses the central helix (22), which is believed to initiate the fusion of viral and host membranes (23). These conserved regions may offer reasonable drug targets because they are more likely to avoid the evolution of drug resistance.

### Evidence of adaptation at multiple genome positions

The observation of parallel evolution in independent lineages enables us to pinpoint specific genome positions that likely increase the fitness of SARS-CoV-2 in the human host. The extreme 5’ and 3’ ends of the genome contained the highest concentration of parallel substitutions (Fig. 2a). Despite their high frequencies, these substitutions were observed exclusively in genomes originating from the same laboratory, which suggests they are sequencing errors rather than authentic mutations. Therefore, we chose to focus on substitutions found in genomes from at least four of the 529 contributing laboratories to mitigate the presence of lab-specific sequencing errors.

We observed two substitutions with more than 30 cases of parallelism across SARS-CoV-2 genomes (Fig. 2b). The most frequent substitution occurred 50 different times at position 11,083, which results in a non-synonymous change (L37F) in nsp6, a transmembrane protein localized to the endoplasmic reticulum and implicated in formation of autophagosomes (24–27). The substitution at 11,083 occurred nearby another frequent substitution at position 11,074 that is synonymous. Both substitutions were conversions to uracil at sites adjacent to eight consecutive uracils in the genome (Fig. 3), suggesting they may occur more frequently due to an increased mutation rate at homopolymeric sites (28). A similar conversion to uracil at position 21,575 is located in the middle of 7 other uracils and results in a non-synonymous change (L5F) to the protein sequence of S (Fig. 3). Three other substitutions were adjacent to at least 3 uracils in the genome: positions 9,474, 26,681, and 28,253. The high frequency of substitutions next to poly(U) tracts is likely due to increased mutation rates at these positions, although we cannot rule out that they may also have adaptive significance.

**Figure 3.**
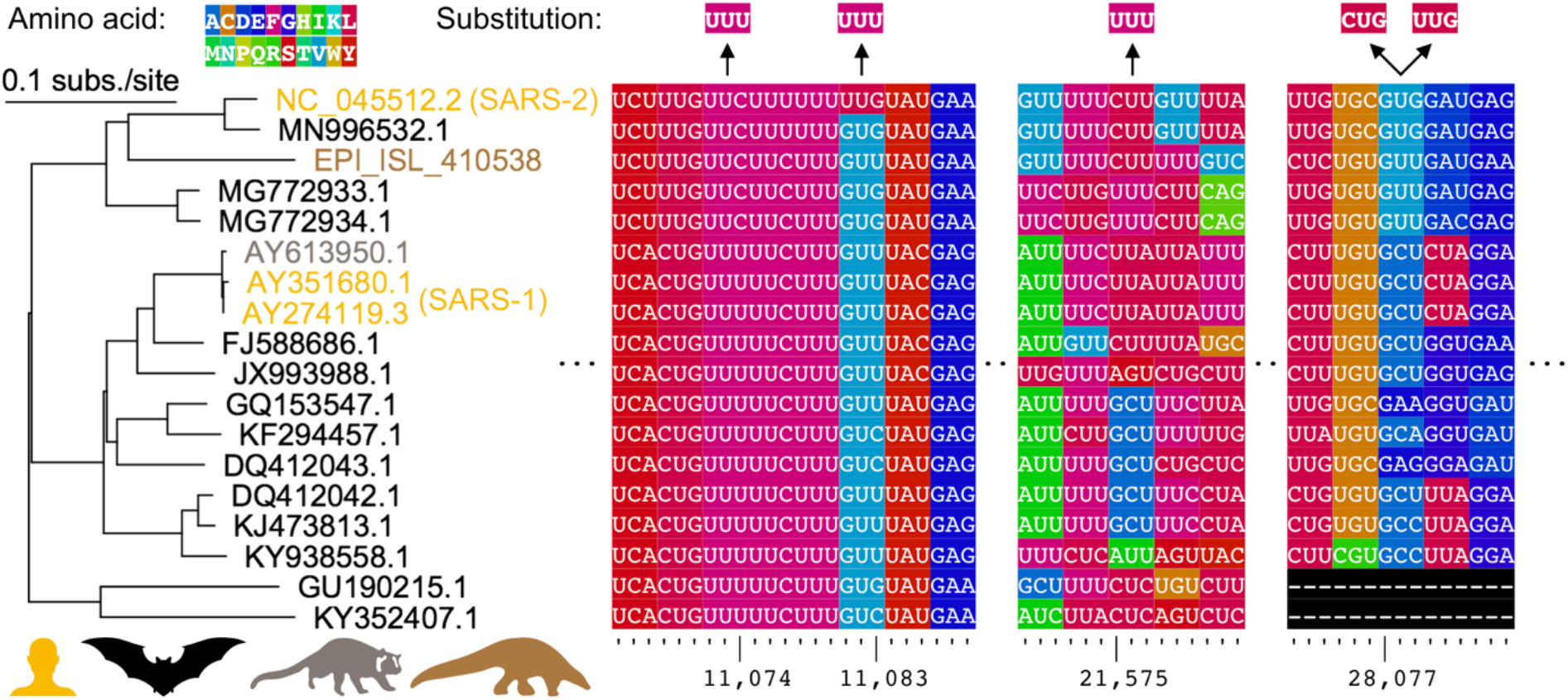
Comparative genomics of parallel substitutions in protein coding regions. A midpoint rooted maximum likelihood phylogenetic tree constructed from SARS-related coronavirus genome sequences with accession numbers colored by host species (human, bat, masked palm civet, or pangolin). Codons are colored by their corresponding amino acid with frequent nucleotide substitutions shown relative to the reference SARS-CoV-2 sequence (top). Poly(U) sequences are found surrounding the substitutions at positions 11,074, 11,083, and 21,575. The two substitutions observed at position 28,077 result in conversions to the same amino acid in ORF8.

The next most frequent substitution occurred 16 times at position 16,887 and results in a synonymous change to nsp3. There is presently no evidence that this mutation is involved in RNA base pairing, and it is located in a region of the genome with relatively little conserved RNA secondary structure (29). The most frequent non-C-to-U substitution was A10323G, which results in a non-synonymous change (K90R) to the protease nsp5. This amino acid replacement is distally located from the active site and the nsp5 dimer interface (30), suggesting it may not be of adaptive significance. We observed a similar substitution, A21137G, which results in a non-synonymous change (K160R) to nsp16. However, this residue is distant from the active site and from the nsp16/nsp10 interface (31), suggesting its replacement could be of little consequence.

Two different nonsynonymous mutations in the N gene, encoding the nucleocapsid protein, repeatedly evolved in a disordered linker domain between structural capsid elements (32). These mutations, R185C and T205I, alter a region acting as an RNA chaperone that facilitates template switching and RNA synthesis during replication (33, 34). Similarly intriguing, we observed divergent nucleotide substitutions, G28077C/U, that both result in the same non-synonymous change (V62L) to ORF8. This region of ORF8 is missing in some SARS-related viruses (Fig. 3), and underwent repeated deletions during the SARS-CoV epidemic (35). ORF8 is known to rapidly evolve and the necessity of its role in the human host remains contentious (36).

We sought to determine whether any of these eighteen highly parallel mutations (Fig. 2b) could be attributable to common sequencing errors. We reasoned that sequencing errors would be randomly distributed across the phylogenetic tree, whereas adaptive mutations are likely to expand in size along a specific lineage. Therefore, we calculated the probability of finding a mutant clade of size R or larger by chance given each substitution’s observed frequency. For example, the substitution G11074U had a largest clade size of R=4, for which we observed 24 mutants among 12,435 genomes. In this case, the probability of observing four or more adjacent mutants on the phylogenetic tree is much less than 10^-6^. Extremely small p-values were found for all parallel substitutions reported here, except the mutation at 9,474 which was only supported by singletons (R=1).

In this study, we determined that SARS-CoV-2 is evolving predominantly under purifying selection that purges most mutations since they are deleterious. This suggests that SARS-CoV-2 was well-poised to invade the human population, although it continues to adapt to humans through specific mutations that may accumulate in individual genomes as SARS-CoV-2 continues to evolve. The few highly parallel substitutions that we observed offer intriguing avenues for further investigation, as most are cryptic and located in poorly characterized regions of the SARS-CoV-2 genome. Notably, some genes acquired relatively few mutations, which implies strict sequence constraints that may focus drug development strategies against these gene products. The paucity of mutations overall suggests that coronaviruses are well-suited to jumping between hosts and caution should be taken to avoid direct or indirect contact with their animal reservoirs. This is further corroborated by the relatively small number of genome positions that have undergone multiple parallel substitutions despite a plentiful supply of mutations.

## METHODS

### Genome collection and comparison

Complete (> 29,000 nucleotide) SARS-CoV-2 genomes were downloaded from GISAID (37) on May 2^nd^, 2020. Genomes with more than 500 degeneracies (e.g., N’s) were removed, resulting in a collection of 12,435 genomes, of which 12,285 had a known date of collection that we used as a proxy for the duration of growth relative to the first genome (2019-12-24). Genomes were aligned to the SARS-CoV-2 reference genome (NC_045512.2) using the DECIPHER (v2.16.1) (38, 39) package for the R (v3.6.1) programming language (40). Genomic distance was defined as the number of positions differing from the reference genome without considering insertions or deletions, which were very infrequent.

To create Figure 4, viruses closely related to SARS-CoV-2 were selected from a recent study (1) and supplemented with one sequence derived from a pangolin host in another study (2). Genomes were aligned and a maximum likelihood tree was created using DECIPHER with the best fitting evolutionary model.

### Identification of parallel substitutions across independent lineages

Starting from the set of all SARS-CoV-2 genomes, we constructed a multiple sequence alignment, matrix of pairwise nucleotide identity, and rooted neighbor joining tree using DECIPHER. Sequences were compared at each site in the reference sequence to identify independent substitutions on the phylogenetic tree. That is, we mapped mutations onto tips of the phylogenetic tree and propagated them back till they coalesced at a common ancestor (edge). This enabled us to count the number of independent substitutions that were inherited by one or more strains. To increase robustness to the tree topology, we ignored single reversions to the ancestral character that occurred within a clade sharing a derived character.

This process resulted in an integer representing the number of parallel substitutions occurring at each position in the reference sequence. We determined that eight or more independent substitutions was statistically significant for C to U transitions (p < 0.001, Poisson distribution and Bonferroni correction) given the observed mutations rates and assuming mutations are randomly distributed along the genome. All other substitutions (e.g., G to U) required fewer cases of parallelism to achieve the same degree of statistical significance.

Conserved regions were defined as stretches (≥ 100 nucleotides) of the genome where the average number of independent substitutions fell below 0.2 (Fig. 2a). To improve the identification of conserved regions, we applied a center-point moving average function that smoothed the mutation signal across the genome. A similar process was used to determine the bias in synonymous versus non-synonymous substitutions within protein coding regions (Fig. 2a). A fully reproducible and open source analysis pipeline is provided on GitHub (https://github.com/digitalwright/ncov).

## ACKNOWLEDGEMENTS

We are grateful for the contributions of laboratories involved in collecting samples and creating the dataset made available by GISAID.

